# Genomic selection in organisms with biphasic lifecycles: a *Saccharina latissima* (sugar kelp) case study

**DOI:** 10.1101/2022.08.01.502376

**Authors:** Mao Huang, Kelly R Robbins, Yaoguang Li, Schery Umanzor, Michael Marty-Rivera, David Bailey, Margaret Aydlett, Jeremy Schmutz, Jane Grimwood, Charles Yarish, Scott Lindell, Jean-Luc Jannink

## Abstract

Sugar kelp (*Saccharina latissima*) has a biphasic life cycle, allowing selection on both the diploid sporophytes (SPs) and haploid gametophytes (GPs). We trained a genomic selection (GS) model from farm-tested SP phenotypic data and used a mixed-ploidy additive relationship matrix to predict GP breeding values. Top-ranked GPs were used to make crosses for further farm evaluation. The relationship matrix included 866 individuals: a) founder SPs sampled from the wild; b) progeny GPs from founders; c) Farm-tested SPs crossed from b); and d) progeny GPs from farm-tested SPs. The complete pedigree-based relationship matrix was estimated for all individuals. A subset of founder SPs (*n* = 58) and GPs (*n* = 276) were genotyped with Diversity Array Technology and whole genome sequencing, respectively. We evaluated GS prediction accuracy via cross validation on farm-tested SPs in two years using a basic GBLUP model. We also estimated the general combining ability (GCA) and specific combining ability (SCA) variances of parental GPs. A total of 11 yield-related and morphology traits were evaluated. The cross validation accuracies for dry weight per meter (*r* ranged from 0.16 to 0.35) and wet weight per meter (*r* ranged 0.19 to 0.35) were comparable to GS accuracy for yield traits in terrestrial crops. For morphology traits, cross validation accuracy exceeded 0.18 in all scenarios except for blade thickness in the second year. Accuracy in a third validation year for dry weight per meter over a confirmation set of 87 individuals was 0.31.

## Introduction

Sugar kelp, *Saccharina latissima*, is a brown seaweed and a winter crop that is economically and ecologically important in the eastern North Pacific and the North Atlantic Oceans (Augyte *et al*. 2017; Kim *et al*. 2017, 2019; Yarish *et al*. 2017; Mao *et al*. 2020; Umanzor *et al*. 2021). Sugar kelp contains nutritional compounds, such as antioxidants, minerals, and vitamins, and has been primarily grown for human consumption. The demand for other uses, such as animal feed, cosmetics, alginates, fertilizers, and biofuels, is increasing rapidly (Rey *et al*. 2019; Kirkholt *et al*. 2019; Vijn *et al*. 2020). As a future potential feedstock for generating biofuels, sugar kelp has advantages over land-based crops due to its high polysaccharide content and its cultivation that requires no land, fresh water, or fertilizer (Kerrison *et al*. 2015; Marinho *et al*. 2015; Lüning and Mortensen 2015; Duran-Frontera 2017; Bruhn *et al*. 2019; Deng *et al*. 2020). The global demand for sugar kelp biomass is increasing and driving the expansion of kelp farming and cultivation programs in the U.S., with similar programs initiated in Europe (Kim *et al*. 2015, 2019; Marinho *et al*. 2015; Augyte *et al*. 2017; Yarish *et al*. 2017). In the United States, kelp farming is emerging as a sustainable mariculture activity providing new economic opportunities and revitalizing waterfronts.

Sugar kelp, like other kelp species, has a biphasic life cycle. Adult sporophytes (SPs) produce sori that release meiospores which develop into female or male haploid gametophytes (GPs). Once GPs reach fertility, they mate to form the next generation of diploid juvenile SPs that further grow into mature SPs (Umanzor *et al*. 2021; Huang *et al*. 2022). This life cycle allows for selection on both GP and SP phases within one breeding cycle (Peteiro *et al*. 2016; Huang *et al*. 2022). Genomic selection is a breeding tool that builds a prediction model using a set of individuals (the training population, TP) with known phenotypic data and genotypic data to predict the performance of individuals with unknown phenotypic data (prediction population, PP) (Meuwissen *et al*. 2001). The relatedness of individuals in TP and PP can be calculated using a common set of markers or pedigree information. As genotyping technologies advance and their cost decreases over time, GS could improve the selection gain per unit of time and cost compared to conventional phenotypic selection in breeding programs (Jannink *et al*. 2010). However, this tool has not been evaluated in any kelp breeding program, to the best of our knowledge. The GS predictive approach can be especially useful for assisting the selection of individuals that are difficult or near impossible to phenotype, like the GP lifestage. Here we report the first GS prediction study in *Saccharina latissima*, for both kelp yield and morphological traits.

Genomic selection has been evaluated as a breeding tool for more than a decade in different terrestrial crops, such as in maize, soybean, wheat, rice, sorghum, and many others (Zhao *et al*. 2012; Jarquín *et al*. 2014; Rutkoski *et al*. 2015; Huang *et al*. 2018, 2019; Fernandes *et al*. 2018). The accuracy of GS, defined as the correlation of phenotypically estimated values and the genomic estimated breeding values determines the usefulness of GS in a program (Rabier *et al*. 2016). The GS accuracies for yield traits in previous studies varied from essentially zero to 0.75 (Zhao *et al*. 2012; Dawson *et al*. 2013; Fernandes *et al*. 2018; Stewart-Brown *et al*. 2019). Several factors are known to affect GS prediction accuracies, including the number of markers used (Zhong *et al*. 2009; Asoro *et al*. 2011), the size of the TP (Asoro *et al*. 2011), the relatedness of individuals between TP and PP (Clark *et al*. 2012; Sallam *et al*. 2015), and the extent of linkage disequilibrium between the markers and causal loci (Zhong *et al*. 2009; Brito *et al*. 2011). Aside from those factors, different statistical models give different prediction accuracies (Heslot *et al*. 2012).

We are interested in evaluating the accuracy of GS in sugar kelp, and we considered several model options. A basic genomic Best Linear Unbiased Prediction (GBLUP) model often provides adequate accuracies when compared to other models including Bayesian approaches (Heslot *et al*. 2012; Sallam *et al*. 2015; Huang *et al*. 2016). These models can be extended to account for genotype by environment interaction (GxE) effects which affect selection accuracy between environments for both GS and phenotypic selection (Resende and Muñoz Del Valle 2011; Lado *et al*. 2016). Models incorporating parental information with general combining ability (GCA) and specific combining ability (SCA) effects account for non-additive gene action and have been reported to be beneficial in millet breeding programs (Jarquin *et al*. 2020).

Our objective was to evaluate the accuracy of GS in a sugar kelp breeding program for both yield and morphological traits in the context of kelp’s biphasic life cycle. To obtain these accuracies, we modified standard genomic relationship matrices used in GBLUP models to include both haploid and diploid individuals. Different GS models were assessed, including the basic GBLUP model and a model with GCA and SCA components. We evaluated accuracies within two training years using cross validation and predicted a third validation year’s data. The SPs evaluated in the third year were made from crosses chosen based on haploid gametophyte breeding values.

## Materials and Methods

### Population

The complete study population was comprised of 866 unique individuals, which included founder SPs sampled from the wild in 2018 (*n* = 104, Mao et al., 2020), GPs derived from the founders (*n* = 439), SPs from experimental crossing of GPs, that were evaluated on farm in 2019 and 2020 (*n* = 245), and GPs derived from the 2019 farm-tested SPs (*n* = 78). A total of 248 experimental and repeated reference crosses were included across years: 124 in Yr2019 and 129 in Yr2020, with five experimental crosses in common between Yr2019 and Yr2020. In addition, a confirmation population was created in Fall 2020 by crossing GPs to produce SPs that were evaluated in Yr2021. The confirmation population produced *n* = 87 plots with useful data.

### Genotyping

The founder SPs samples were genotyped for single nucleotide polymorphisms (SNPs) using the DArTSeq platform by Diversity Array Technology LLC, as described in Mao et al. ((Mao *et al*. 2020)). A subset of 4,906 markers was retained based on minor allele frequency greater than 5% and fewer than 5% missing values (Mao *et al*. 2020). A total of 58 genotyped founder SPs contributed to downstream members of the population. Gametophyte DNA was extracted using the Macherey-Nagel NucleoSpin Plant II Maxi Kit (Macherey-Nagel, Düren, Germany) with a modified protocol. In brief, 24 mg (fresh weight) of gametophyte culture was transferred from Erlenmeyer flasks into 1.5 mL centrifuge tubes. The tubes were centrifuged at 21,000 rcf for 2 min using an Eppendorf centrifuge 5424. The supernatant was removed. The tubes containing the gametophytic biomass were capped and submerged in liquid nitrogen for 20 seconds. The frozen samples were then ground up manually for 30 seconds using a plastic pestle. Once samples were ground, the CTAB extraction buffer with repeated wash steps protocol was followed. The DNA was whole-genome sequenced at the HudsonAlpha Institute. Kelp DNA was cleaned using a DNAeasy PowerClean Pro Cleanup kit (Qiagen) and amplified Illumina libraries were generated in 96 well format using an Illumina TruSeq nano HT library kit using standard protocols. Sequencing was performed on a Illumina NovaSeq 6000 instrument at 2x150 base pair read length. Raw reads are available at the NCBI Short Read Archive, Accession ZYXnnnnnnnn. Sequence reads from 278 GPs (all generated from 2018 founder SPs) were aligned to a reference genome (A publication describing this genome is in preparation) using BWA (Li and Durbin 2010). The average read depth across GPs ranged from 4 to 37. Downstream sequence data formatting, SNP variant calling and filtering were done using SAMtools (Li *et al*. 2009), Picard tools in java (*http://broadinstitute.github.io/picard/*), BCFtools (Li 2011) and VCFtools (Danecek *et al*. 2011). A total of 909,747 bi-allelic SNP markers with good quality were retained by removing markers with more than 20% missing values and minor allele frequency less than 5%. These markers were used to evaluate the population structure among GPs via Principal Component Analysis (PCA).

### Mixed-ploidy additive relationship matrix

We recorded the full pedigree connecting all individuals (*n* = 866), both SPs and GPs. Using this pedigree we calculated a coefficient of coancestry matrix (CCM) across all individuals. This CCM tracks *haplotypes* so that each SP is represented by two rows and two columns in the matrix, and each GP is represented by one row and one column. A simple tabular method is used (Emik and Terrill 1949). All diagonal elements of the matrix are equal to 1, because each haplotype has a probability of 1 of being IBD with itself. For founder SPs all off-diagonal values are set to zero, reflecting the assumption that founder SPs were non-inbred and unrelated to each other. The two rows of a diploid SP, offspring of two haploid GPs, are copies of these parental GP rows, because each GP contributes its exact genome as half of the diploid SP genotype. The row of a haploid GP, offspring of a diploid SP, is the mean of the two rows representing its parent SP, because random Mendelian segregation suggests that each GP has a one-half probability of inheriting one or the other of the SP haplotypes. In our case, these rules led to a CCM that had 1215 rows and columns (2x104 founder SPs + 439 first generation GPs + 2×245 first generation SPs + 78 second generation GPs = 1215).

The rules for converting the CCM of identical by descent probabilities into an additive relationship matrix are straightforward to derive. Consider the additive effect of GP1, *A*_*GP*1_, carrying allele *i*, the allele substitution effect of which is *α*_*i*_. Then, *A*_*GP*1_ = *α*_*i*_. The additive relationship between GP1 and GP2, which carries allele *i′* is

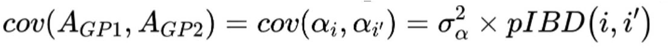

where 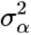 is the variance of allele substitution effects and *pIBD*(*i, i*′) is the probability that alleles *i* and *i*′ are identical by descent, as given by the CCM. Similarly, the additive relationship between GPl and SPl, which carries alleles *i*′and *j*′ is

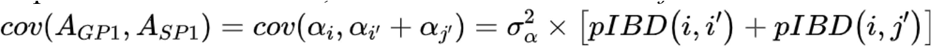

Finally, the additive relationship between two SPs is the sum of the four pairwise IBD probabilities between their respective alleles. Consequently, the CCM can be “condensed” into a mixed-ploidy additive relationship matrix as follows. Relationships between pairs of GPs are represented by single cells and are unchanged. Relationships between a GP and an SP are represented by two cells which are summed to obtain the single additive relationship between them. Relationships between two SPs are represented by four cells which are also summed to obtain the single additive relationship between them. Note that in standard diploid quantitative genetics, the constant of proportionality commonly used to relate the additive relationship matrix to the additive covariance matrix among individuals is the additive genetic variance, which is two times the variance of allele substitution effects as defined above 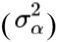. Given that we had both haploid and diploid individuals, we found it easier to work with 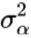 as the constant of proportionality. A consequence of this choice is that the diploid narrow-sense heritability is calculated as 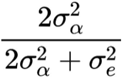, where 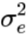 is the error variance on farm-tested SPs.

### Mixed-ploidy combined pedigree and marker relationship matrix

For historical reasons, two marker systems were used: one on the founder SPs (DArTSeq) and one on derived GPs (whole-genome sequencing). Markers were imputed within the genotyped SP founders and within the GP subset using the EM algorithm in the rrBLUP package in R (R Core Team, 2022). Marker-based additive relationships between haploid GPs were calculated using the same formula as the *A*.*mat()* function in the rrBLUP package (Eq. 15, (Endelman and Jannink 2012)) except that the marker dosage matrix has dosages of 0 and 1 prior to centering, and the coefficient of 2 is removed from the denominator. Marker-based additive relationships between founder SPs were calculated using the *A*.*mat()* function of the rrBLUP package (Endelman 2011). For this matrix to be appropriately scaled relative to the GP matrix, it was multiplied by 2 (equivalent to removing the coefficient of2 from the denominator of the GP matrix).

Following these calculations, three matrices were available: 1) a pedigree-based matrix including all individuals, 2) a marker-based matrix for the founder SPs, and 3) a marker-based matrix for derived GPs. These three relationship matrices were combined using the *CovCombR* package (Akdemir *et al*. 2020), with marker-based matrices weighted twice as heavily as the pedigree-based matrix. The Wishart EM-algorithm was used to estimate the combined relationship matrix from partial samples (Akdemir *et al*. 2020). We denote this relationship matrix **G** below.

### Phenotyping

The Yr2019 trial had a shorter growing season relative to the Yr2020 and Yr202 l trials. For Yr2019, outplanting occured Jan. 26th, 2019 and harvest May 28th, 2019 (Umanzor *et al*. 2021). For Yr2020 and Yr2021, outplanting occurred respectively on Dec. 6th, 2019 and Dec. 9th 2020, and harvest on June 8th 2020 and June 7th 2021 (Li et al. 2022). We measured eleven traits: Wet Weight per Meter (WWpM, kg), percent Dry Weight (pDW, %), Dry Weight per Meter (DWpM, kg), Ash-Free Dry Weight per Meter (AshFDWpM, kg), percent Ash content (Ash,%), and Blade Density (BD, number of blades/m), Blade Length (BL, cm), Blade maximum Width (Bm Wid, cm), Blade Thickness (BTh, mm), Stipe Length (SL, cm), and Stipe Diameter (SDia, mm). Percent Dry weight at the plot level was derived from subsample measurements. Detailed experimental design and trait measurements were reported in Umanzor et al. ((Umanzor *et al*. 2021)) and Li et al. (2022). Briefly, an augmented block design was used where farmed lines with sequential plots were laid out in parallel, and plots were grouped in blocks across lines. Different GPs were used as reference check SPs on the farm in Year2019 and Year2020 due to limitations in obtaining sufficient biomass of the same GPs for making reference checks in the second year. Due to differential survival of the SPs, we observed heterogeneity of rope coverage within plots. To minimize the impact of this heterogeneity, we removed data from plots where < 10% of the plot rope was covered by the SP. All phenotypic data were natural log transformed to normalize the data and stabilize error variances for all analyses.

### Heritability estimation and same trait genetic correlation between years

The relationship matrix **G** was used to estimate the additive genetic variance. Similar to Atanda et al. (Atanda *et al*. 2021), we fit a univariate heterogeneous variance model using the ASReml-R package to fit genotype within environment (environment being year in our case) effects (Butler *et al*. 2017), with an unstructured variance-covariance matrix, **G**_0_, across two years. The diagonal elements of **G**_0_ are the additive genetic variance 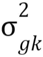 within the *k*^*th*^ year, and off diagonal elements are the genetic covariance between years:

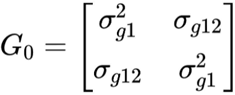

The analysis was done for each trait. For simplicity in constructing the design matrix, the line, block and year effects were treated as fixed, whereas genotypes were treated as random. The formula was:

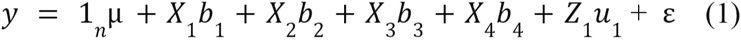

where y is the vector of phenotypes with length *n*, and *n* is the total number of observations across two years; 1_*n*_ is the vector of 1s and μ is the overall mean; *b*_1_ is the fixed effects of experimental versus check crosses; *b*_2_ is the fixed effects the two years, *b*_3_ is the fixed effects of growth lines nested within year, *b*_4_ is the fixed effects of blocks nested within year, and *X*_1_ to *X*_4_ are the incidence matrices for fixed effects. Vector *u*_1_ is the random effects of genotype within year, and *Z*_1_ is the incidence matrix for *u*_1_; finally ε is the error term. The random effect is distributed as *u*_1_~N[0, **(G**_0_**⊗G)]**, where ⊗ is the Kronecker product, and **G**_0_ and **G** are as described above.

The error was distributed as ε ~N(0, **R):**

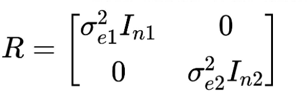

where 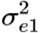 and 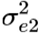 are within-year error variances and *I*_n1_ and *I*_n2_ are identity matrices of the size of the number of observations within years. The narrow sense heritability was estimated within the *k*^*th*^ year as:

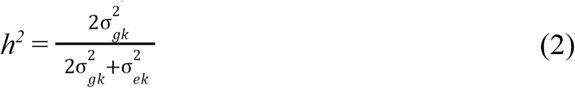

The output from model (1) were GBLUPs estimated for each year. These GBLUPs are breeding values (BVs). We also obtained combined BVs across years by averaging the Yr2019 and Yr2020 GBLUPs. The genetic correlation for the same trait across Yr2019 and Yr2020 was computed using the variance-covariance components estimated from this model (Table 1). The correlation coefficients among BVs for all traits using BLUPs averaged across years were calculated. A histogram of the combined BVs for SP plots from Yr2019 and Yr2020 was plotted (Supp. Fig. 1).

**Table 1.**
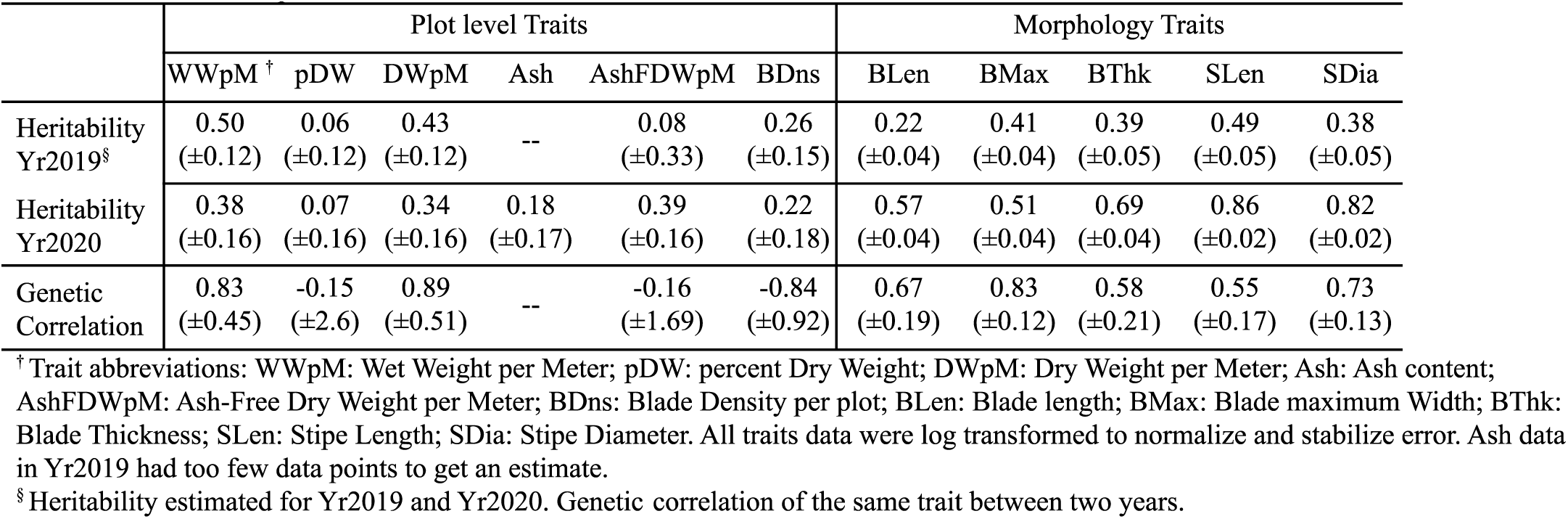
Heritability across two years and genetic correlation between years for ten different traits. Values in parenthesis are approximate standard errors for variance components obtained in *ASReml-R*.

|  | Plot level Traits |  |  |  |  |  | Morphology Traits |  |  |  |  |
| --- | --- | --- | --- | --- | --- | --- | --- | --- | --- | --- | --- |
|  | WWpM <sup>†</sup> | pDW | DWpM | Ash | AshFDWpM | BDns | BLen | BMax | BThk | SLen | SDia |
| Heritability Yr2019<sup>§</sup> | 0.50<br>(±0.12) | 0.06<br>(±0.12) | 0.43<br>(±0.12) | -- | 0.08<br>(±0.33) | 0.26<br>(±0.15) | 0.22<br>(±0.04) | 0.41<br>(±0.04) | 0.39<br>(±0.05) | 0.49<br>(±0.05) | 0.38<br>(±0.05) |
| Heritability Yr2020 | 0.38<br>(±0.16) | 0.07<br>(±0.16) | 0.34<br>(±0.16) | 0.18<br>(±0.17) | 0.39<br>(±0.16) | 0.22<br>(±0.18) | 0.57<br>(±0.04) | 0.51<br>(±0.04) | 0.69<br>(±0.04) | 0.86<br>(±0.02) | 0.82<br>(±0.02) |
| Genetic Correlation | 0.83<br>(±0.45) | -0.15<br>(±2.6) | 0.89<br>(±0.51) | -- | -0.16<br>(±1.69) | -0.84<br>(±0.92) | 0.67<br>(±0.19) | 0.83<br>(±0.12) | 0.58<br>(±0.21) | 0.55<br>(±0.17) | 0.73<br>(±0.13) |

<sup>†</sup> Trait abbreviations: WWpM: Wet Weight per Meter; pDW: percent Dry Weight; DWpM: Dry Weight per Meter; Ash: Ash content; AshFDWpM: Ash-Free Dry Weight per Meter; BDns: Blade Density per plot; BLen: Blade length; BMax: Blade maximum Width; BThk: Blade Thickness; SLen: Stipe Length; SDia: Stipe Diameter. All traits data were log transformed to normalize and stabilize error. Ash data in Yr2019 had too few data points to get an estimate.
<sup>§</sup> Heritability estimated for Yr2019 and Yr2020. Genetic correlation of the same trait between two years.

### Genetic correlation between dry weight per meter and morphology traits

The genetic correlation among traits was estimated within each year due to the large GxE for most traits. We used a multivariate model in the ASReml-R package (Butler *et al*. 2017). Due to model convergence issues, traits were analyzed in pairs and only for dry weight per meter and five individual morphological traits. An unstructured variance-covariance matrix across all traits was assumed in this model (Jia and Jannink 2012; Fernandes *et al*. 2018):

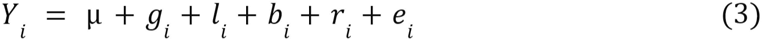

Where *Y*_*i*_ is the vector of phenotypic observations for the *i*^*th*^ genotype: *Y*_*i*_ = [*Y*_*i*1_*Y*_*i*2_ … *Y*_*ip*_],and *p* is the *p*^*th*^ trait,μ = [μ_1_ μ_2_… μ_p_],where is μ is the mean for the *p*^*th*^ trait, and *l*_*i*_, *b*_*i*_ and *r*_*i*_. are the line, block and year effects for the *i*^*th*^ individual, respectively. The *g*_*i*_ and *e*_*i*_ terms are the genetic effects and residual effects for the *i*^*th*^ individual, with *g*_*i*_ = [*g*_*i*1_ *g*_i2_ … *g*_*ip*_] and *e*_*i*_ = [*e*_*i*1_ *e*_*i*2_ … *e*_*ip*_]. Across all individuals, the vectorized genetic effects were distributed as **g** ~ *N*_*np*_ (0, **G**_0_ ⊗ **G)**, where **G**_0_ is the among-trait genetic variance-covariance matrix and **G** is the additive relationship matrix. Similarly, **e** ~ *N*_*np*_(0, **R**_0_⊗**I**), where **R**_0_ is the among-trait residual variance-covariance matrix and I is an identity matrix showing our assumption that residuals are uncorrelated across plots (Fernandes *et al*. 2018).

### Genomic prediction between diploid and haploid life stages

The breeding values for all 866 SPs from Yr2019 and Yr2020, and their parental GPs were obtained from the GBLUP model. Haploid GPs were selected to make crosses to create the confirmation population of diploid SPs grown in Yr2021. These GPs were used for crossing based on their available biomass (which determines the number of crosses to which they can contribute) and the following crossing design criteria: first, the GPs were ranked based on their dry weight per meter GEBVs. Parental GPs’ were then selected if their BVs were top ranked (*n* = 42), bottom ranked (*n* = 3), and randomly ranked (*n* = 25). In addition, we included GPs whose parental SPs were sampled from otherwise unrepresented locations (*n* = 5). Finally, we generated crosses that were repeats from the previous year (*n* = 12).

The GEBVs of Yr2021 diploid SPs were obtained as the sum of GEBVs from their parental female and male GPs, as estimated from equation (1), and their phenotypic observations were estimated using the Best Linear Unbiased Estimator (BLUEs), as explained below.

We assessed GS prediction accuracies as the correlation between BVs and BLUEs for all Yr2021 SPs (Table 3), for SPs within the category of parental GPs being top ranked, for SPs in the randomly selected category, and for SPs that were repeats from previous year (Supplemental Table 1). For the 12 common SPs between Yr2020 and Yr2021, we also estimated the phenotypic correlations between their within-year BLUEs. The analysis was done for each trait and prediction accuracy for all available traits that were recorded, even though selection of the Yr2021 population was only based on dry weight per meter.

### Within-year cross validation genomic prediction accuracy

For modeling simplicity, GS model prediction accuracy comparisons were done with a two-step approach using the Yr2019 and Yr2020 data. The experimental block and line effects were statistically controlled for within each year to obtain BLUEs of SPs in *ASReml-R*. These BLUEs were then used as the response values to evaluate prediction accuracy under four different GS models using the BGLR package in R. The BLUEs were first estimated using the following model:

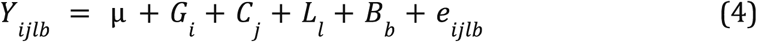

where *Y*_*ijlb*_ is the *ijlb*^*th*^ observation, μ is the overall mean, *G*_*i*_ is the *i*^*th*^ genotype, *C*_*j*_ is the *j*^th^ check group that distinguishes between experimental and check crosses, *L*_*l*_ is the *l*^*th*^ line, and *B*_*b*_ is the *b*^*th*^ block, and *e*_*ijlb*_ is the error associated with the *ijlb*^*th*^ observation. The BLUEs for Yr2021 were estimated using the same model, but without a group effect *C*_*j*_ because none of the reference check plots generated high quality data.

Four GS models were run with a 10-fold cross validation scheme within each year. Each randomization scheme was repeated 20 times. The four models were:

#### A) General combining ability (GCA) and specific combining ability (SCA) using combined pedigree and marker relationship matrix

The mixed-ploidy combined pedigree and marker relationship matrix was used. General combining ability (GCA) and specific combining ability (SCA) components for the GP parents were included in the model with a formula adapted from Jarquin et al. ((Jarquin *et al*. 2020)):

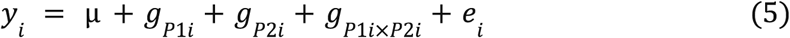

where *y*_*i*_ is the BLUE of the *i*^*th*^ individual, *g*_*P*1*i*_ and *g*_*P*2*i*_ are the genetic effects of parent GPl and parent GP2. And *g*_*P*1*i*×*P*2*i*_ is the interaction effect of two parents, *e*_*i*_ is the g_*P1ixP2i*_ residual effect. The GCA and SCA account for the genetic main effects and the interaction effects, respectively, where the SCA variance-covariance matrix is the cell-wise product of the elements in variance-covariance matrix from parent 1 and from parent 2 (Jarquin *et al*. 2020).

For the purpose of estimating variance components and their 95% credible intervals (CIs), model (5) was also analyzed without cross validation (all data were used to predict BVs of all individuals). Variance components estimated were for female parent GCA (GCA_FG), male parent GCA (GCA_MG), SCA and error (varE). This model was run within each year and each running process was repeated 20 times. The lower- and upper-bound values of the 95% CI were averaged across the 20 replicates (Table 2b).

**Table 2.** Genetic correlation among traits using a multi-trait model in ASReml-R. Analysis was done within years. The genetic correlation values for Yr2019 are in the upper diagonal and for Yr2020 are in the lower diagonal.

| Yr2019 |  |  |  |  |  |  |
| --- | --- | --- | --- | --- | --- | --- |
| Yr2020 | DWpM<sup>†</sup> | BLen | BMax | BThk | SLen | SDia |
| DWpM | -- | 0.45 | -0.40 | 0.73 | 0.53 | 0.34 |
| BLen | 0.88 | -- | 0.02 | 0.90 | 0.08 | 0.11 |
| BMax | 0.15 | -0.39 | -- | -0.90 | 0.05 | 0.49 |
| BThk | 0.82 | 0.69 | 0.23 | -- | 0.22 | 0.10 |
| SLen | 0.77 | 0.64 | 0.68 | 0.58 | -- | 0.84 |
| SDia | 0.78 | 0.65 | 0.77 | 0.69 | 0.95 | -- |

<sup>†</sup> For trait abbreviations refer to Table 1.

#### B) GCA+SCA model using only pedigree based relationship matrix

The same GCA +SCA model as formula (5) was used, except that the additive relationship matrix was estimated based only on pedigree. The GS 10-fold cross validation accuracies were obtained as in *A*.

#### C) Genomic Best Linear Unbiased Predictor (GBLUP) model using combined pedigree and markers relationship matrix

A basic GBLUP mixed model was used:

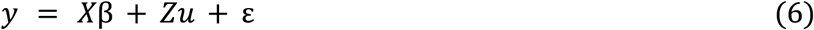

where X and Z is the design matrix for fixed effects β and for random effects u, respectively. ε is the error and 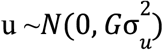, and **G** is the relationship matrix using combined pedigree and markers relationship matrix as estimated above. The 10-fold cross validation scheme being used was the same as the other models.

#### D) GBLUP model using only pedigree based relationship matrix

The same model as in formula (6) was used except that the **G** matrix was replaced by a relationship matrix estimated using pedigree information only.

## Results

### Population structure of GPs

Two clear subpopulations were revealed, separating GPs from GOM and SNE locations (Figure 1a). This subpopulation structure is consistent with the founder SP subpopulation structure (Mao *et al*. 2020). No strong subpopulation structure was observed within the GOM region (Figure 1b).

**Figure 1.**
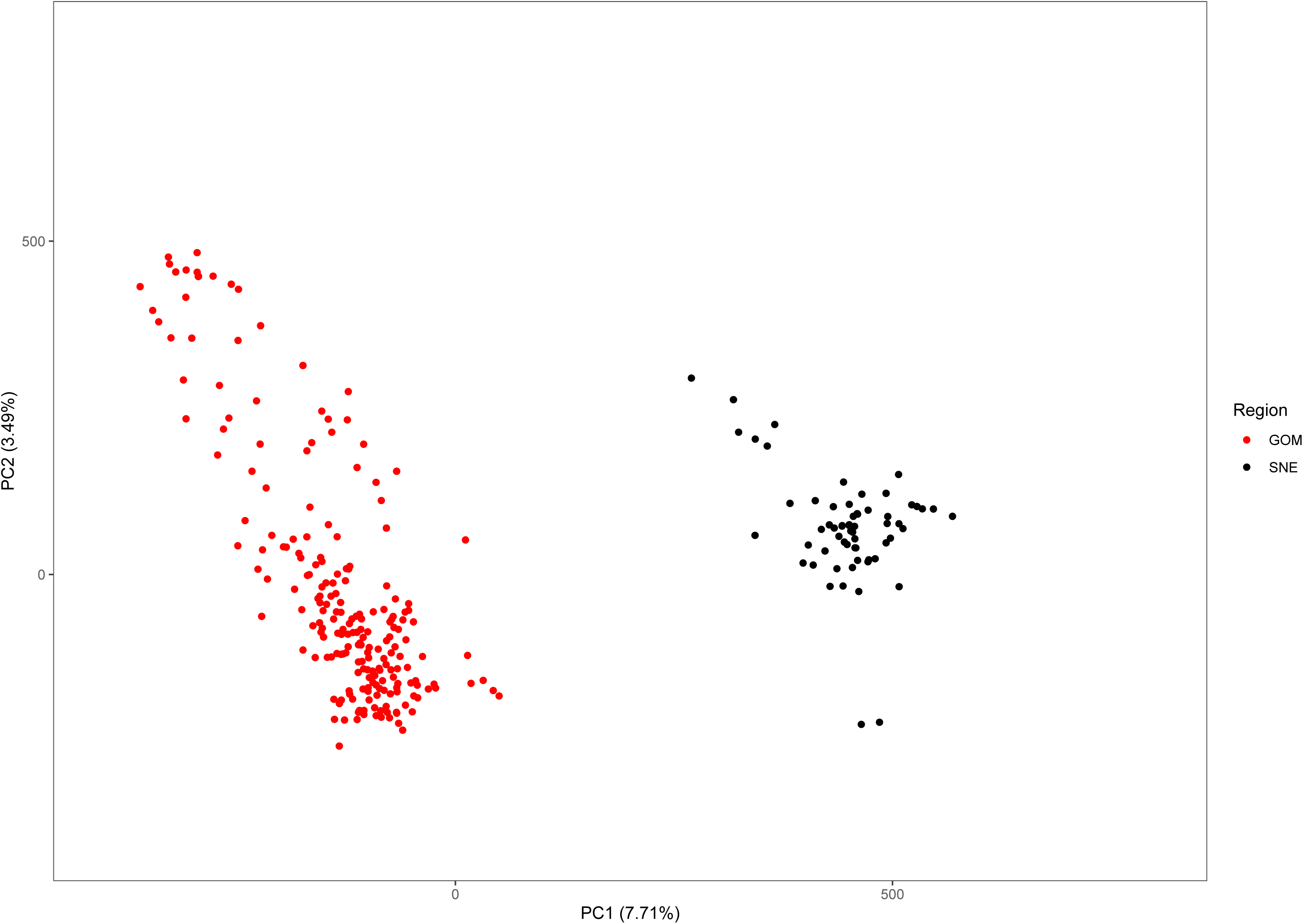
a) Principal Component Analysis (PCA) for 276 GPs using 909,749 SNPs. Red symbols: Gulf of Maine (GOM), black symbols: Southern New England (SNE). b) PCA for GPs within the GOM region. Gametophyte parent SPs were collected from these locations: CB: Casco bay farm, CC: Cape Cod, Check: gametophytes came from checks crosses, JS: Fort Stark, LD: Lubec Dock, LL: Lubec Light, NC: Newcastle, NL: Nubble Light, OD: Outer Dock, 01: Orr’s Island, and SF: Sullivan Falls.

### Heritability and genetic correlation between years

Trait heritabilities using two years’ data with the heterogeneous error model ranged from 0.06 to 0.86 (Table 1). Among six plot level (yield-related) traits, the highest heritability was for wet weight per meter in (0.50 in Yr2019), and lowest was for percent dry weight (0.06 in Yr2019). Among five morphology traits within each year, the highest heritability was for stipe length (0.86 in Yr2020), and the lowest was for blade length (0.22 in Yr2019, Table 1). All morphology traits had higher heritability in Yr2020 compared to Yr2019. In Yr2020, the heritabilities for morphology traits, measured at the individual blade level, were higher than plot-level yield-related traits (Table 1). The genetic correlation for all traits between the two years ranged from negative (ash related traits, percent dry weight and blade density) to strongly positive, with the highest being dry weight per meter (0.89 ± 0.51, Table 1). The heritability of ash free dry weight per meter was low one year (0.08, Yr2019), but higher the next year (0.39, Yr2020) while the between-year genetic correlation was nominally negative (−0.16 ± 1.7). The standard errors associated with between-year genetic correlation estimates were large, especially for traits with negative correlations, including percent dry weight (−0.15 ± 2.6) and blade density (−0.84 ± 0.92). None of the genetic correlations with nominally negative values were significantly different from zero. Lack of genotype by environment interaction results in a genetic correlation of 1 (Cooper and DeLacy 1994). Even though the genetic correlations with negative nominal estimates were not significantly different from zero, they indicated large genotype by year interactions.

In the multivariate analysis to estimate genetic correlations between traits, we only included the results between dry weight per meter and the morphological traits due to model convergence problems. Given the genotype by environment interaction effects, the genetic correlation analysis was done within years. Dry weight per meter had higher genetic correlation with morphological traits in Yr2020 than in Yr2019. It was not genetically correlated with blade maximum width in either year (*r* = −0.40 in Yr2019 and *r* = 0.15 in Yr2020, Table 2). For individually measured morphological traits, blade length and blade thickness were strongly correlated in both years (*r* = 0.90 in Yr2019 and *r* = 0.69 in Yr2020, Table 2). Stipe length and stipe diameter had the highest consistent correlations in both years (*r* = 0.84 in Yr2019 and *r* = 0.95 in Yr2020, Table 2). Among-trait correlations of breeding values (BVs) averaged across two years showed that dry weight per meter was moderately correlated with most morphology traits (*r* = 0.47 for Blade Thickness *tor=* 0.58 Blade Length), though it was uncorrelated with blade maximum width (*r* = −0.11, Figure 2). The correlation of Blade density BV was relatively strongly correlated with Wet and Dry weight per meter (*r* = 0.55 and *r* = 0.54, respectively, Figure 2).

**Figure 2.**
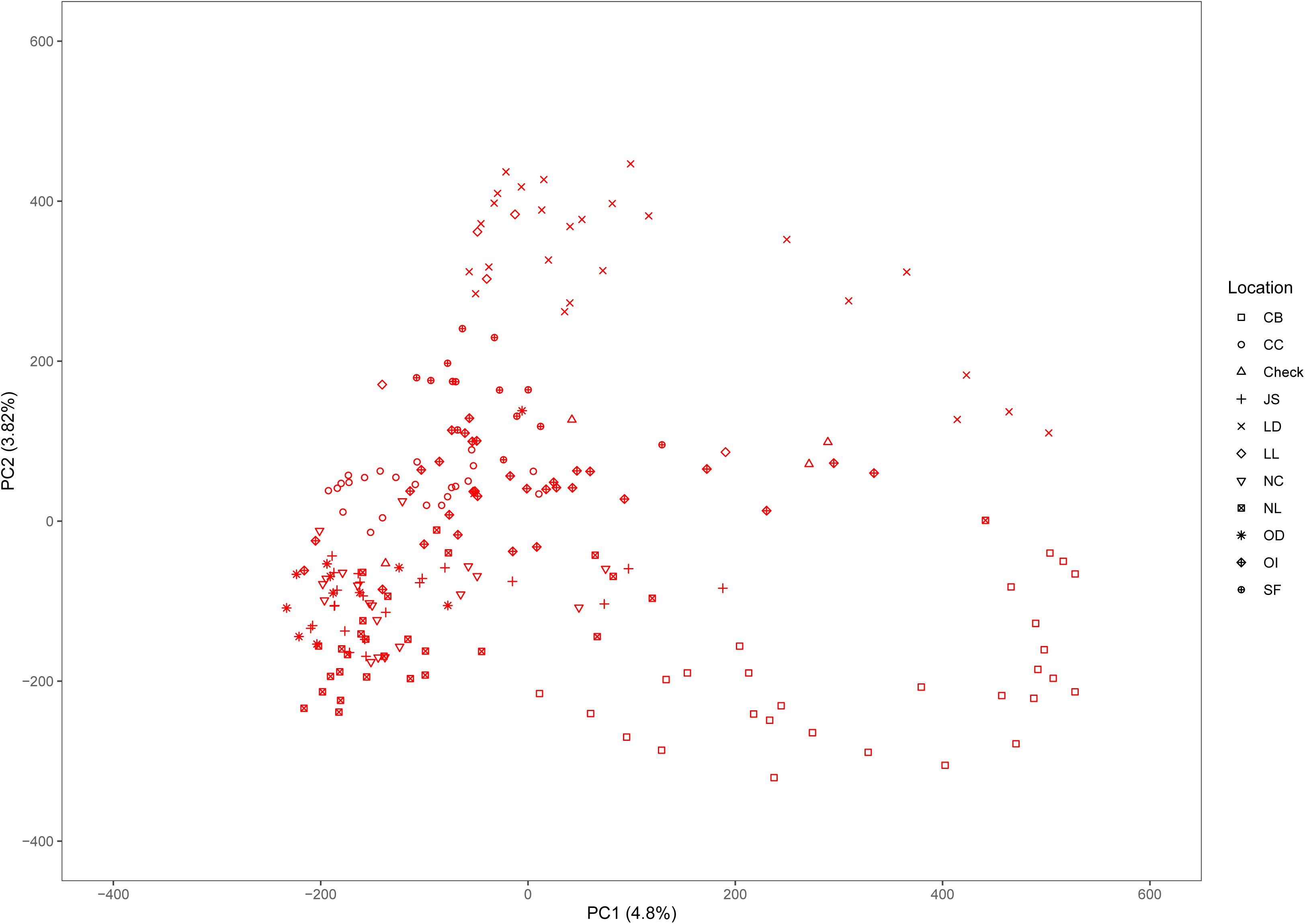
Correlation plot of SP Breeding values estimated using both Yr2019 and Yr2020 data with the heterogeneous error variance GBLUP model. The combined BVs are the mean of estimates for Yr2019 and Yr2020.

### Within-year cross validation genomic prediction accuracies

Genomic prediction accuracy with the GCA+SCA model using both pedigree and markers for all traits via 10-fold cross validation ranged from negative for percent dry weight, blade thickness to 0.48 for stipe length and stipe diameter (Table 3). For yield traits, the maximum prediction accuracy was for both wet weight and dry weight per meter (0.32 to 0.35, Table 3). Averaged across traits, prediction accuracy within Yr2020 was slightly lower numerically than within Yr2019 (Table 3). This lower accuracy also occurred across morphological traits despite their higher heritability in Yr2020 than Yr2019.

**Table 3.** Genomic prediction accuracy from 10-fold cross validation within each year. Each analysis was run with 20 replicates.

| Model | TP<sup>†</sup> | PP | WWpM<sub>§</sub> | pDW | DWpM | Ash | AshFD<br>WpM | BDns | BLen | BMax | BThk | SLen | SDia |
| --- | --- | --- | --- | --- | --- | --- | --- | --- | --- | --- | --- | --- | --- |
| GCA+SCA<br>(Pedigree<br>& Markers) | Yr2019 | Yr2019 | 0.35 | -0.01 | 0.35 | 0.28 | 0.20 | 0.24 | 0.38 | 0.48 | 0.29 | 0.48 | 0.35 |
|  | Yr2020 | Yr2020 | 0.34 | 0.02 | 0.32 | 0.08 | 0.31 | 0.12 | 0.32 | 0.31 | -0.19 | 0.38 | 0.47 |
| GCA+SCA<br>(Pedigree<br>only) | Yr2019 | Yr2019 | 0.31 | -0.06 | 0.32 | 0.23 | 0.13 | 0.24 | 0.31 | 0.45 | 0.23 | 0.43 | 0.28 |
|  | Yr2020 | Yr2020 | 0.28 | 0.00 | 0.25 | 0.06 | 0.25 | 0.10 | 0.23 | 0.30 | -0.18 | 0.33 | 0.44 |
| GBLUP<br>(Pedigree<br>& Markers) | Yr2019 | Yr2019 | 0.34 | -0.02 | 0.33 | 0.23 | 0.12 | 0.17 | 0.42 | 0.55 | 0.20 | 0.50 | 0.36 |
|  | Yr2020 | Yr2020 | 0.22 | 0.04 | 0.22 | 0.04 | 0.26 | 0.09 | 0.33 | 0.28 | -0.23 | 0.30 | 0.34 |
| GBLUP<br>(Pedigree<br>only) | Yr2019 | Yr2019 | 0.32 | -0.04 | 0.33 | 0.15 | 0.02 | 0.19 | 0.32 | 0.51 | 0.18 | 0.44 | 0.29 |
|  | Yr2020 | Yr2020 | 0.19 | 0.04 | 0.16 | 0.00 | 0.20 | 0.10 | 0.26 | 0.26 | -0.18 | 0.25 | 0.30 |

<sup>†</sup>TP: Training Population; PP: Prediction Population.
<sup>§</sup> For trait abbreviations refer to Table 1.

The GS cross validation accuracies from the GCA+SCA model using both pedigree and markers across all traits in two years was slightly higher (*r* = 0.27, averaged first two rows in Table 3) than that from the model of GCA+SCA with pedigree only (*r* = 0.22) and the base GBLUP model with pedigree and markers (*r* = 0.23) or GBLUP model with pedigree only (*r* = 0.20). While these differences were not significant for any single trait comparison, there was only one case (Blade thickness in Yr2020) when prediction accuracy was higher without than with marker information (Table 3). The trend for GS within-year prediction accuracy was similar to that for heritability values across traits in that morphological traits tended to have higher GS prediction accuracy (and higher heritability) than yield-related traits (Table 1 and Table 3).

We compared the variance within each year due to female parents (GCA_FG) with those due to male parents (GCA_MG) using the MCMC-derived samples from the posterior distributions of these parameters. Across all yield-related and morphology traits, GCA_FG was greater than GCA_MG in 15 out of 22 trait-year combinations (Table 4). However, GCA_FG was significantly greater than GCA_MG only for stipe diameter in Yr2020, as determined by the fact that GCA_FG was greater than GCA_MG in over 97.5% of posterior samples (Table 4). Variance component results also explain the relative superiority of the GCA+SCA model over the base GBLUP model: in 17 out of 22 trait-year combinations the SCA variance component was greater than the average of the female and male GCA variance components (Table 4).

**Table 4.** The 95% credible interval of posterior distribution for variance components estimates: variance due to GCA components from Female GPs (GCA_FG) and from Male GPs (GCA_MG), SCA component, and Error Variance (varE) based on the Variance component estimates. Model used within year BLUEs as input and the marker and pedigree based combined relationship matrix in BGLR.

|  | Yr2019 |  |  |  | Yr2020 |  |  |  |
| --- | --- | --- | --- | --- | --- | --- | --- | --- |
|  | GCA_FG | GCA_MG | SCA | varE | GCA_FG | GCA_MG | SCA | varE |
| WWP<sup>†</sup> | 0.01 to 0.08 | 0.01 to 0.06 | 0.02 to 0.11 | 0.05 to 0.15 | 0.05 to 0.32 | 0.03 to 0.14 | 0.05 to 0.31 | 0.14 to 0.44 |
| pDW | 0.00 to 0.01 | 0.00 to 0.01 | 0.01 to 0.03 | 0.01 to 0.04 | 0.00 to 0.02 | 0.00 to 0.02 | 0.01 to 0.04 | 0.03 to 0.07 |
| DWpM | 0.00 to 0.003 | 0.00 to 0.03 | 0.00 to 0.05 | 0.00 to 0.01 | 0.01 to 0.04 | 0.00 to 0.02 | 0.01 to 0.05 | 0.02 to 0.07 |
| Ash | 0.00 to 0.005 | 0.00 to 0.02 | 0.00 to 0.01 | 0.00 to 0.01 | 0.00 to 0.01 | 0.00 to 0.04 | 0.00 to 0.01 | 0.00 to 0.01 |
| AshFDWpM | 0.00 to 0.003 | 0.00 to 0.00 | 0.00 to 0.03 | 0.00 to 0.002 | 0.00 to 0.02 | 0.00 to 0.01 | 0.00 to 0.02 | 0.01 to 0.03 |
| BDns | 0.04 to 0.36 | 0.03 to 0.15 | 0.07 to 0.45 | 0.22 to 0.66 | 0.03 to 0.21 | 0.03 to 0.20 | 0.06 to 0.38 | 0.17 to 0.53 |
| BLen | 0.01 to 0.04 | 0.01 to 0.07 | 0.01 to 0.08 | 0.04 to 0.11 | 0.02 to 0.11 | 0.01 to 0.07 | 0.02 to 0.13 | 0.07 to 0.20 |
| BMax | 0.01 to 0.06 | 0.01 to 0.05 | 0.01 to 0.05 | 0.02 to 0.07 | 0.01 to 0.04 | 0.01 to 0.07 | 0.01 to 0.06 | 0.03 to 0.10 |
| BThk | 0.00 to 0.002 | 0.00 to 0.03 | 0.00 to 0.01 | 0.00 to 0.01 | 0.00 to 0.04 | 0.00 to 0.04 | 0.00 to 0.01 | 0.00 to 0.01 |
| SLen | 0.02 to 0.10 | 0.01 to 0.09 | 0.02 to 0.13 | 0.06 to 0.17 | 0.06 to 0.34 | 0.03 to 0.13 | 0.05 to 0.27 | 0.12 to 0.38 |
| SDia | 0.00 to 0.02 | 0.00 to 0.01 | 0.00 to 0.03 | 0.01 to 0.03 | **0.02 to 0.08\*** | 0.01 to 0.03 | 0.01 to 0.06 | 0.02 to 0.07 |

<sup>†</sup> For trait abbreviations refer to Table 1.
\* Bolded: For over 97.5% of posterior samples GCA\_FG > GCA\_MG

### Genomic prediction in both diploids and haploids

Due to the large genotype by environmental interactions and the differences in error variance observed between Yr2019 and Yr2020,lwe verified the ability of phenotypes from these years to predict performance in Yr2021. Our main interest was the BV for dry weight per meter (DWpM) trait for which we selected GPs to make crosses. GS prediction accuracy for Yr2021 SPs was defined as the correlation coefficient between trait BLUEs observed in 2021 and their predicted BVs, which were calculated as the sum of their parental FG and MG BVs. For the DWpM trait, GS accuracy for the 87 crosses in Yr2021 was *r* = 0.30 (using Yr2019 BVs) and *r* = 0.31 (using Yr2020 or combined BVs, Table 5). Over all other traits, GS accuracy ranged from negative (ash related traits, percent dry weight and blade density) to 0.80 (stipe length with Yr2019 GEBVs, Table 5). When heritability was high for a trait within a year, the prediction accuracy using that year’s data to predict Yr2021 also tended to be high (Tables 1 and 5). Breeding values for Founder SPs, GPs generated from founders, and SP progenies from those GPs all centered on zero (Fig. 3). In contrast, GPs generated from SPs that were evaluated and selected on farms and SP progenies from those selected GPs deviated markedly from zero (Fig. 3).

**Table 5.** Genomic prediction accuracy using heterogeneous error variance model, the same model as used in Table la. The training set was Yr2019 and Yr2020 SP data and BVs for haploid GPs were predicted. For the Yr2019+Yr2020 combination set, the BVs for haploid GPs were the mean of their BVs estimated from Yr2019 and Yr2020. The prediction set derived from Yr2021 SPs.

| Model | GPs BVs predicted from | WWpM<sup>†</sup> | pDW | DWpM | Ash | AshF DWpM | BDns | BLen | BMax | BThk | SLen | SDia |
| --- | --- | --- | --- | --- | --- | --- | --- | --- | --- | --- | --- | --- |
| GBLUP<br>(Pedigree<br>&<br>Markers) | Yr2019 | 0.29 | 0.10 | 0.30 | -- | -0.10 | -0.21 | 0.26 | 0.34 | 0.17 | 0.52 | 0.38 |
|  | Yr2020 | 0.30 | 0.02 | 0.31 | 0.04 | 0.19 | 0.22 | 0.29 | 0.36 | 0.11 | 0.44 | 0.30 |
|  | Yr2019+<br>Yr2020 | 0.30 | 0.08 | 0.31 | -- | 0.19 | -0.04 | 0.29 | 0.35 | 0.14 | 0.49 | 0.33 |

<sup>†</sup> For trait abbreviations refer to Table 1. For Ash, the within Yr2019 training sample size was small ( $n=45$ ) and results were not included.

**Figure 3.**
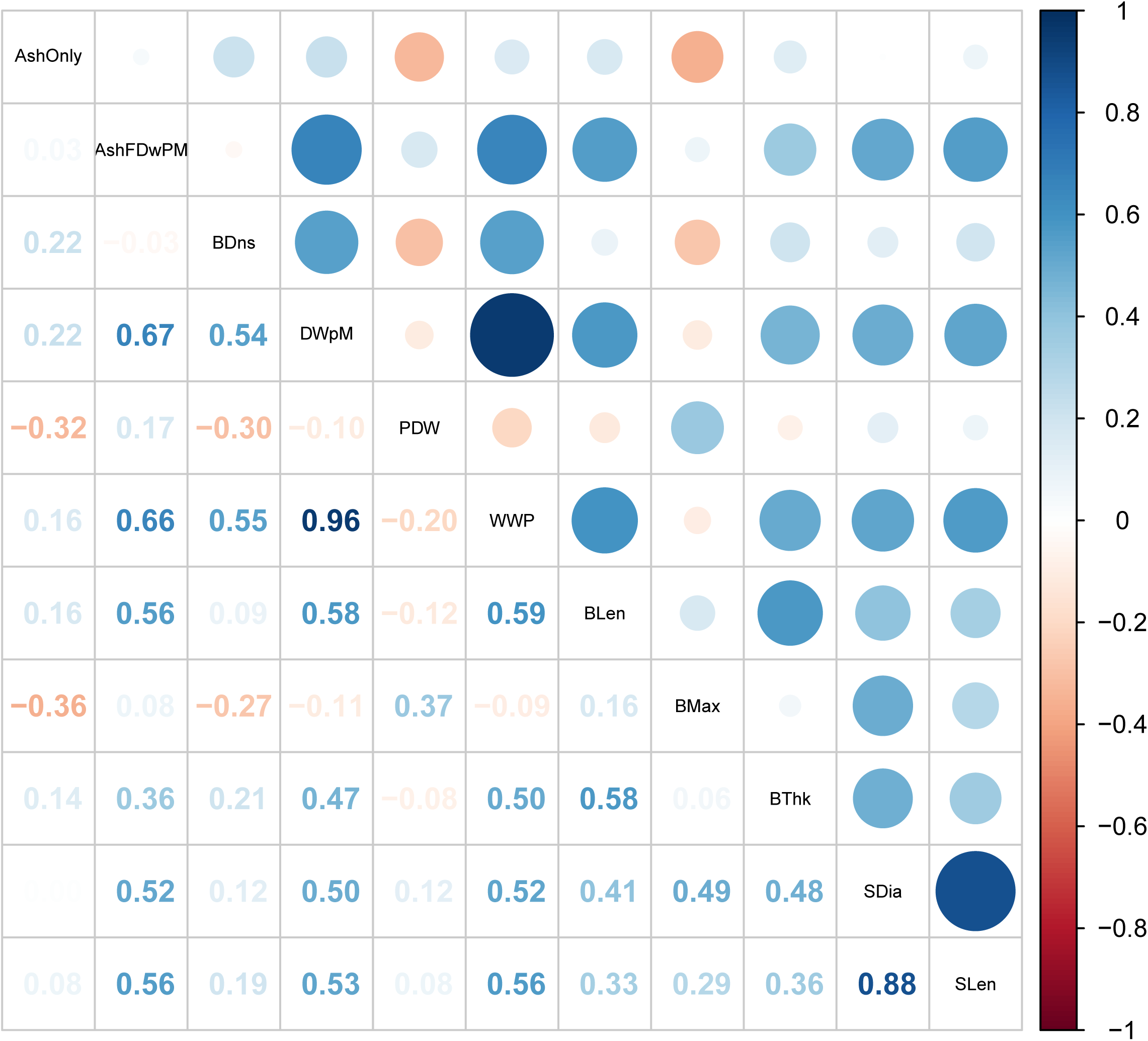
Box plots of estimated breeding values of DWpM (Dry Weight per Meter, log transformed) for 517 GPs and 436 SPs. The SPs included these subcategories: Founder SPs without progeny (these SPs were collected from the wild but did not produce GP progeny, denoted FndSP−), Founder SPs with progeny (FndSP+), SP19_Fnd (SPs tested on farm in Yr2019 generated using GPs from Founders), SP20_Fnd (SPs tested in Yr2020), SP21_Fnd (SPs tested in Yr2021), and SP21_Frm (SPs tested in Yr2021, generated using GPs collected from SPs tested on farm in Yr2019). The GPs included GPs collected from Founders (GP_Fnd) and GPs collected from SPs tested on farm in Yr2019 (GP_Frm).

## Discussion

### Heritability, Genetic Correlation, and Environmental Variation

At this early stage in the development of technologies to phenotype diverse SP genotypes of sugar kelp in farm-like conditions, the heritabilities of yield-related traits are moderate to low (Table 1). The differences we observed in estimated heritability for the yield-related traits between years (Table 1) was likely due to changes in environmental conditions from one year to the next. The Yr2019 trial was planted later and harvested earlier than the Yr2020 trial, reducing total growth potential. Possibly as a result of these differing conditions, we found a negative genetic correlation between the two years for percent dry weight, ash related traits, and blade density, revealing high GxE and large year-to-year effects for these traits. Differences between years can also result from differences in nutrient availability which, in many nearshore sites, depends on runoff caused by rainfall events that vary annually (Grebe *et al*. 2021).

Phenotyping sugar kelp traits is challenging (Umanzor *et al*. 2021). Due to phenotyping limitations and labor constraints (all measurements made on hundreds of crosses within 48 hours of harvest), we relied on subsampled data for blade density and percent dry weight to approximate whole plot traits. In theory each plot consists of uni-clonal individuals such that subsamples should be uniform. Yet due to environmental and possible blade density effects, we observed non-uniform growth across the plot. Lack of uniformity among subsamples contributed to the error variance. To minimize the impact of this heterogeneity, we filtered data based on a visual score of plot uniformity (see Methods). We modified the measurement of dry weight related traits from Yr2019 to Yr2020. In Yr2019, percent dry weight was measured by a single value per plot where subsamples were combined and weighed together. In Yr2020, we measured the percent dry weight for the 10 largest blades out of the subsamples separately. Neither approach gave a good heritability. This low heritability is in contrast to land crops where percent dry weight generally has high heritability (e.g., (Rabbi *et al*. 2020) for root dry weight in cassava), though comparisons to traits in other domesticated species are perhaps unwarranted given the divergence, approximately 1,600 million years ago, of kelps from other eukaryotes (Hedges *et al*. 2004).

Because of the longer growing season in 2020 compared to 2019, dry weight per meter and ash measurements collected in 2020 were markedly higher than those in 2019. To normalize the data, we log transformed the raw data in all analyses leading to a stabilized error variance. However, because connectivity between years came primarily from evaluating related SPs rather than from repeating evaluation of the same SPs across years, the genetic effects could also be partially confounded by year effects in the model.

Morphology traits were directly measured at the individual blade level, and we randomly measured 15 blades/plot in Yr2019. We modified the sampling procedure in Yr2020 by measuring only the largest ten blades from 3 subsamples within each plot. This modification ensured individual samples were more uniform within the same plot in Yr2020, likely reducing the error variance and leading to higher heritabilities for morphology traits in Yr2020 (Table 1). Blade length heritability more than doubled in the second year, and all other morphology traits also had increased heritability in Yr2020. In general, when traits had good heritability in both years, the genetic correlation values between years were also reasonable.

Multi-trait genetic correlation results among dry weight per meter and morphology traits indicate genes underlying some of the morphology traits could play a role in contributing to kelp yield. The improved phenotyping protocol in Yr2020 could have contributed to a higher genetic correlation than those in Yr2019. Interestingly, stipe length and diameter were strongly correlated with each other in both years, indicating that they were controlled by the same group of genes across different environments (Egan *et al*. 1990).

Correlations from estimated BVs combined across two years on all plots revealed that dry weight per meter was correlated to other weight related traits except for percent dry weight (Fig 2). If a plot generates larger wet weight, it will also have a larger dry weight per meter and ash free dry weight per meter values (Fig 2). There was a clear correlation between plot level weights and blade density (*r* of 0.54 and 0.55 with dry and wet weight per meter, respectively, Fig 2). High blade density comes from the ability of the juvenile SP to attach to seed-string in the nursery and survive. This trait was not under selection in the wild, so it is not surprising that variation for this trait exists (Paaby and Rockman 2014). Thus, blade density appears to be an important domestication trait.

The BVs of dry weight per meter were correlated to morphology traits, except blade maximum width (Fig 2). A similar pattern of positive genetic correlation was also observed in the multivariate analysis (Table 2). Previous studies reported that the multivariate model is favored as it allows borrowing information between traits and individuals across environments (Atanda *et al*. 2021). The multivariate genetic correlation results indicate that our selection for dry weight per meter could lead to morphologically longer and thicker blades, longer and larger stipes. The fact that wider blades were not particularly favored indicates that “skinny” individuals (designated *Saccharina angustissima*), originating from Casco Bay may have good adaptation to farms (Augyte *et al*. 2018). Indeed, Li et al (2022) showed that SPs with skinny kelp ancestry out-yielded SPs that had no skinny kelp ancestry. In our experiment, among the 42 Yr2021 top ranked SPs, 14 of them (33%) had a skinny kelp derived GP as a parent or grandparent. For comparison, of the 493 cross attempts made for Yr2019 and Yr2020 combined, 123 (25%) had a skinny kelp derived GP as a parent. A chi-square test showed these proportions were not significantly different.

### Genomic prediction within years via cross validation

In previous studies, the GBLUP GS model produced similar accuracies across various traits compared to other GS models such as Bayesian approaches (Heslot *et al*. 2012; Sallam *et al*. 2015). Similarly, we did not observe significant differences between the model prediction accuracy of GBLUP (whether using pedigree alone, or using pedigree plus marker information) and GCA+SCA (whether using pedigree alone or using pedigree and marker information together). Nevertheless, for both GBLUP and GCA+SCA, addition of marker data improved prediction accuracy over pedigree information alone (Table 4). To our knowledge, we are the first to use the algorithm implemented in the *CovCombR* package (Akdemir *et al*. 2020) for an actual selection experiment. This approach allows the combination of data from different genotyping platforms without cross-platform imputation, therefore potentially simplifying the process. Our successful use of the method provides it with some validation.

While the prediction accuracy for GCA+SCA with pedigree and marker information was not significantly different from the other three models, it was numerically superior. There are two biological reasons why this model might be superior. First, we show suggestive evidence that the female contribution to trait expression is moderately more important than the male contribution. Prior studies have lacked sufficient numbers of crosses to assess this issue (Umanzor *et al*. 2021). Relevant mechanisms could be genomic imprinting (Reik and Walter 2001; Yang *et al*. 2021) or simply the fact that the female GP plays a more important role in controlling kelp holdfast and stipe traits and, thus, attachment to the seed string substrate (Garbary and South 2011). Using the GCA+SCA with pedigree and marker model, we could distinguish between GCA variation contributed from the female versus male side. Across 22 comparisons (11 traits in each of two years, Table 4) the female contribution was significantly greater than the male contribution in only one case (Stipe Diameter in Yr2020, Table 4). However, the female contribution was numerically superior to the male contribution in 15 out of 22 cases when they were different (Table 4). A two-sided binomial test would give a probability of 0.06 of that many cases occurring, though we note that the assumption of independence of the 22 cases is probably violated. While the GCA+SCA model allows the importance of parental contributions to differ, the standard GBLUP model forces them to be equal. Second, we also show evidence that the specific combining ability between the parental genomes is often important (Table 4): in 17 out of 22 cases, the SCA variance was greater than the mean of the two GCA variances. Again, given the lack of independence of these cases, we do not have a statistical test to conclusively show that SCA is greater than the GCAs, but this result does suggest it is biologically important. The SCA is caused by non-additive modes of gene action that the GBLUP model does not capture (Lynch and Walsh 1998), therefore providing another mechanism making the GCA+SCA model superior to the GBLUP model.

GS accuracy estimates from cross validation are usually higher than estimates from independent datasets (Crossa *et al*. 2010; Michel *et al*. 2016; Huang *et al*. 2018). The yield-related prediction accuracies via cross validation (*r* ranged from close to zero for ash free dry weight in the absence of marker data to 0.35 for wet weight with marker data, Table 3) fell within the range of those in previous studies in terrestrial crops (Zhao *et al*. 2012; Dawson *et al*. 2013; Michel *et al*. 2016; Stewart-Brown *et al*. 2019). We confirmed that GS prediction accuracy was related to trait heritability (Dawson *et al*. 2013; Lenz *et al*. 2020): in general traits with high heritability gave good prediction accuracy, with an unexplained exception for blade thickness in Yr2020.

### Genomic prediction between diploids and haploids

One of the biggest merits of applying genomic selection is that it can shorten the time per breeding cycle by directly predicting the breeding values of non-phenotyped individuals. In the case of sugar kelp breeding, the breeding germplasm is maintained in the form of GPs in culture. Because SP yield and composition traits cannot be obtained from GPs directly, genomic prediction is the only option for direct selection on those traits. The historical SP data from Yr2019 and Yr2020 was used to predict BVs of GPs, which were then used as parents for next generation SPs in Yr2021. The GS accuracy (*r* ~ 0.30) for dry weight per meter on the confirmation population was similar to the reports in other studies for grain yield (Zhao *et al*. 2012; Stewart-Brown *et al*. 2019). This reasonable prediction accuracy validated our method for calculating the pedigree-based mixed-ploidy relationship matrix and integrating it with marker-based genomic relationship matrices at the different ploidy levels. Simulation has shown that selection during both GP and SP phases of the kelp life cycle will generate the greatest breeding gain per unit time (Huang *et al*. 2022). This empirical research is the first to report that genomic prediction of haploid breeding values works. The use of GS in biphasic organisms, such as sugar kelp, can help breeders achieve higher efficiency in its breeding.

The Yr2021 season was more similar to Yr2020 than Yr 2019 in terms of the overall length of the season and growth of the kelp plots. We were therefore somewhat surprised that Yr2019 and Yr2020 training data gave equal prediction accuracy of the Yr2021 validation data (Table 5). We show that, as for land crops (de Leon *et al*. 2016), genotype by year interaction is an important source of phenotypic variation (Table 1). Large GxE effects require a larger number of environments with repeated plots in order to properly evaluate genotype performance. Previous studies have confirmed that when more information is shared between environments or when sets of genotypes are observed across environments, the prediction accuracy can be increased (Jarquín *et al*. 2014; Lado *et al*. 2016; Jarquin *et al*. 2020). Future kelp breeding efforts need to take this source of variation into account.

### Breeding value changes over generations of breeding cycles

The previous results show some success in genomic prediction. Our empirical evaluations also show success in genomic selection: GPs derived from SPs that were evaluated on farm in Yr2019 (GP_Frm, Fig 3) were superior to GPs derived from founders (GP_Fnd, Fig 3) and, in turn, SPs derived from those GPs (SP21_Frm) were superior to SPs derived from founder GPs (SP19_Fnd, SP20_Fnd, and SP21_Fnd, Fig. 3). We note that the superiority of farm-derived GPs occurred despite minimal selection pressure on the SP parents of those GPs (Supp. Fig. 1). Improvements in our experimental procedures meant that we collected twice as many SPs that had successful spore release in Yr2020 (Supp. Fig. 1, green dashed lines) than in Yr2019 (red dashed lines). This improvement was mainly due to the fact that more plots in Yr2020 than Yr2019 were mature (longer growing season) and hence more plots produced sorus tissue. The Yr2020 plots also had a larger proportion of SPs that were top ranked compared to Yr2019 (Supp. Fig. 1). Greater success in collection of SPs with spore release will lead to higher selection intensity in both the SP and GP phases (Huang et al. 2022). The response of GPs despite low selection pressure in Yr2019 (Fig. 3) suggests that, in addition to artificial selection, some amount of natural selection is also taking place within our breeding program as we shift kelp adapted to attaching to rock substrates to being adapted to the seed string and rope substrates. Continued selection in this way may lead to appreciably “domesticated” kelp in the sense that our germplasm may be better adapted to the off-shore farm environment than to the natural rocky intertidal habitat.

### Future research areas

The use of a pedigree and marker estimated relationship matrix between lab-grown GP cultures and the field-grown SPs enabled us to predict breeding values of the GPs and to select the top ranking ones to make new crosses. Our results indicate that progress can be made in sugar kelp breeding by genomic selection, especially for dry weight per meter. To the best of our knowledge, our study is the first to evaluate the use of GS in kelp breeding, including a mixed-ploidy relationship matrix and the integration of separate genomic relationship matrices at the two ploidy levels. Our results specifically show that wet weight and dry weight per meter can be effectively selected via GS. However, continued efforts in improving nursery/planting and phenotyping methods, as well as increasing the number of plots to be evaluated on farms will be critical for us to continuously improve prediction accuracy. The GS model should also be updated and retrained as we move forward using data from related individuals. The low across-year genetic correlations we observed were concerning (Table 1). These findings need to be backed up with further experimentation, including the addition of common individuals across environments. Finally, our results with the GCA+SCA model suggest that it may be superior to the standard GBLUP model for kelp genomic predictions, but validation of that hypothesis will require more experimental data.

## Data Availability

R Scripts and data sets are available on Github: https://github.com/MaoHuang2020/SugarKelpBreeding/tree/2YrData/TraitAnalyses201003

Raw phenotypic and genotypic data are available on sugarkelpbase.org.

Farm evaluation phenotypic data are at: https://sugarkelpbase.org/breeders/trial/392 https://sugarkelpbase.org/breeders/trial/391 https://sugarkelpbase.org/breeders/trial/388

Genotypic data on the founder SPs are at: https://sugarkelpbase.org/breeders_toolbox/protocol/4

Genotypic data from whole genome sequencing GPs are at: https://sugarkelpbase.org/breeders_toolbox/protocol/5

## Acknowledgments

We acknowledge funding support from the U.S. Department of Energy ARPA-E (DE-AR0000915), and the Massachusetts Clean Energy Center (AmplifyMass). We thank Jeff Glaubitz from Cornell Institute of Biotechnology for consultation on bioinformatics. This work has been published as a reprint in bioRxiv (bioRxiv Locator).

## Author Contributions

MH performed the analyses, wrote the manuscript draft, and revised the manuscript together with other coauthors. KRR and J-LJ guided the analyses. J-LJ contributed to analysis scripts. YL, SU, SL, and CY edited the manuscript. MH, YL, SU, MMR, CY, DB, MA and SL collected phenotypic data. JS and JG performed DNA extraction and sequencing. SL, CY, and J-LJ led the project and all authors read and approved the manuscript.

## Supplemental Figure legends

**Supplemental Figure 1.**
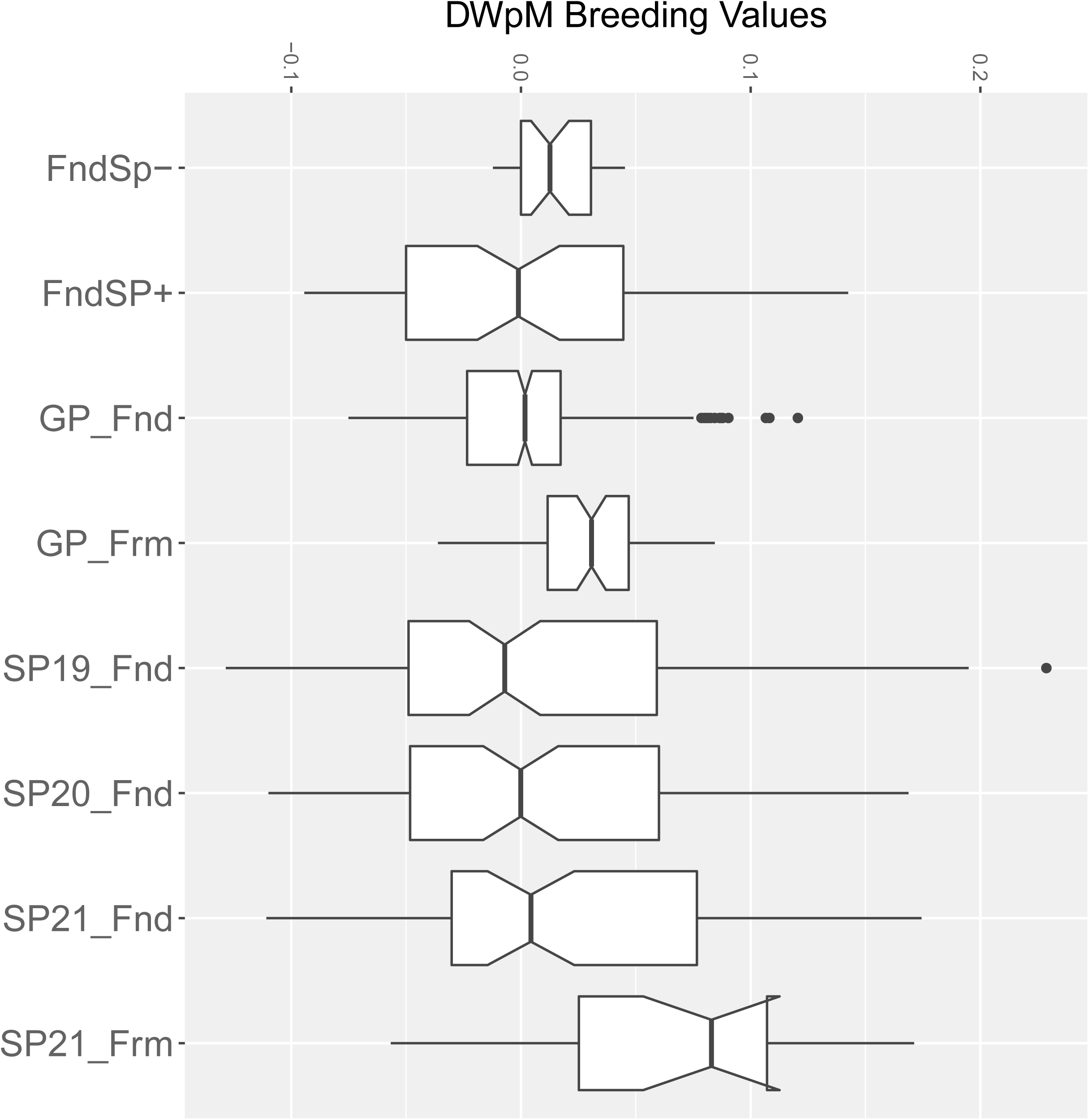
Histogram of estimated breeding values for Dry Weight per Meter, log transformed (DWpM, kg/m) for SPs from Yr2019 and Yr2020. Red dashed lines indicate the Yr2019 plots that had a successful meiospore release leading to isolation of viable GPs which were used for making the 2020-2021 crosses. The green dashed lines indicate Yr2020 plots that had a successful meiospore release.

## Supplemental Tables

**Supplemental Table 1.** Genomic prediction accuracy using heterogeneous error variance structure model to predict performance of Yr2021. The model was a GBLUP model using both pedigree and markers combined relationship matrix, the same model as used in Table 3. The training set was Yr2019 and Yr2020 diploid SPs data, and BVs for haploid GPs were predicted. For the Yr2019+Yr2020 combination set, the BVs for haploid GPs were the mean of their BVs estimated from Yr2019 and Yr2020. The prediction set was Yr2021 SPs, which their estimated BVs were the sum of parental GPs BVs. The GS prediction accuracy is the correlation of assembled BVs for Yr2021 SPs and their BLUEs.

| Category | BVs predicted from | WWpM<sub>s</sub> | pD<sub>W</sub> | DWpM | Ash | AshF DWpM | BD | Blen | BmWid | BTh | SL | SDia |
| --- | --- | --- | --- | --- | --- | --- | --- | --- | --- | --- | --- | --- |
| Top Ranked (n=43) | Yr2019 | 0.20 | 0.14 | 0.24 | -0.02 | -0.04 | -0.18 | 0.28 | 0.37 | 0.24 | 0.41 | 0.40 |
|  | Yr2020 | 0.17 | -0.04 | 0.25 | 0.02 | 0.27 | 0.16 | 0.26 | 0.40 | 0.14 | 0.33 | 0.25 |
|  | Yr2019 +Yr2020 | 0.18 | 0.06 | 0.25 | -0.01 | 0.27 | -0.19 | 0.28 | 0.39 | 0.19 | 0.38 | 0.30 |
| Random Ranked (n=25) | Yr2019 | 0.05 | 0.23 | 0.07 | 0.04 | -0.28 | 0.06 | 0.07 | 0.55 | 0.15 | 0.57 | 0.24 |
|  | Yr2020 | 0.11 | 0.44 | 0.10 | 0.00 | 0.27 | -0.06 | 0.20 | 0.59 | 0.05 | 0.63 | 0.19 |
|  | Yr2019 +Yr2020 | 0.09 | 0.49 | 0.09 | 0.07 | 0.27 | 0.05 | 0.17 | 0.58 | 0.10 | 0.63 | 0.21 |
| Common plots (n=12) | Yr2019 | 0.45 | 0.42 | 0.43 | -0.48 | 0.13 | -0.50 | 0.20 | 0.33 | 0.21 | 0.80 | 0.53 |
|  | Yr2020 | 0.51 | -0.44 | 0.45 | 0.47 | -0.32 | 0.51 | 0.37 | 0.38 | 0.42 | 0.57 | 0.51 |
|  | Yr2019 +Yr2020 | 0.49 | -0.39 | 0.45 | -0.48 | -0.32 | 0.13 | 0.31 | 0.36 | 0.38 | 0.67 | 0.52 |
| Common plots (n=12) | Phenotypic correlation | 0.25 | -0.10 | 0.12 | 0.11 | -0.61 | 0.38 | 0.03 | 0.41 | -- | 0.64 | 0.51 |

<sup>†</sup> For trait abbreviations refer to Table 1a. For phenotypic correlation, blade thickness was not reported as Yr2020 only had 2 data points.

---

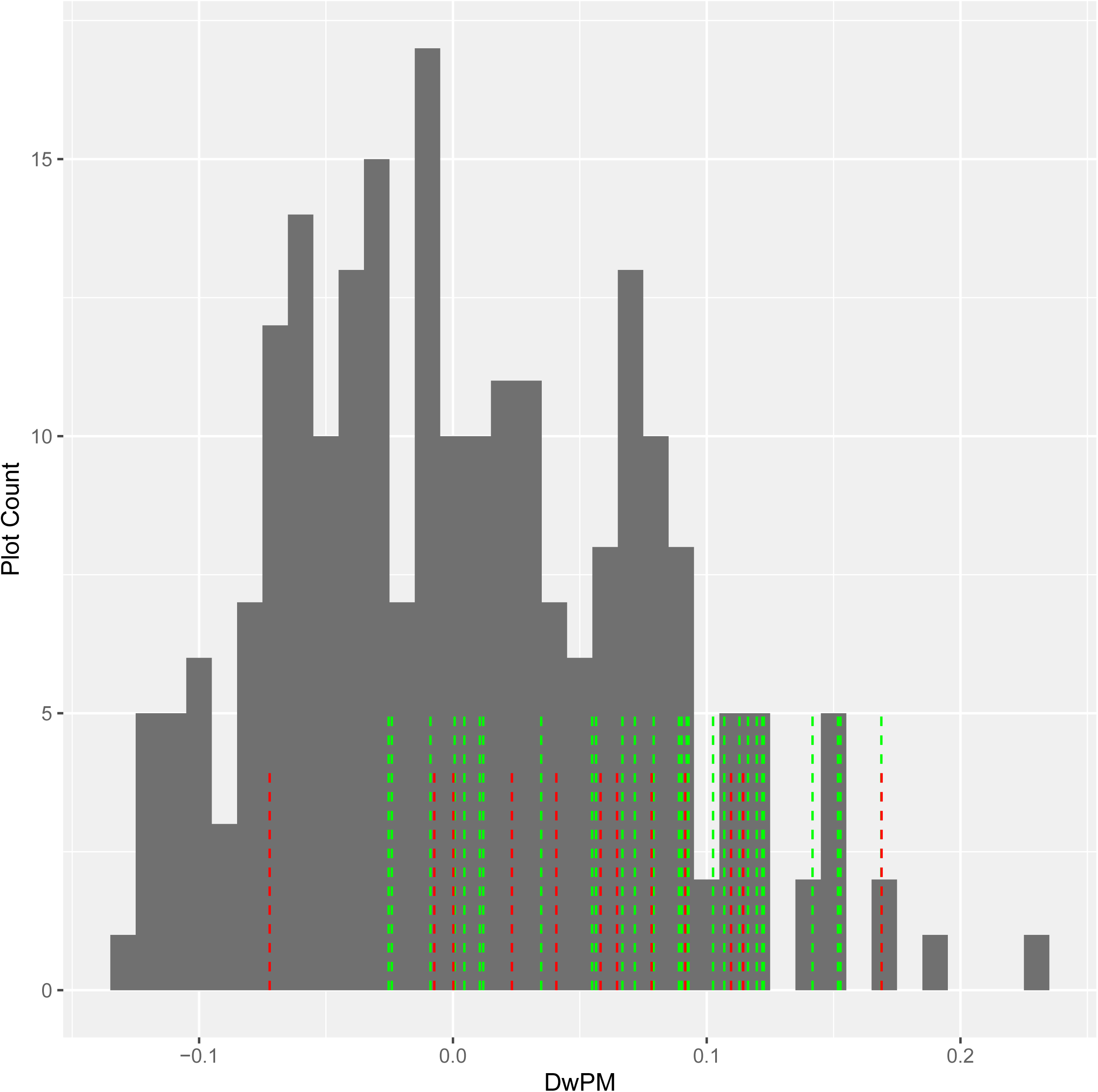

